# Type-2 CD8^+^ T lymphocytes responsive to PGD_2_ and LTE_4_ in severe eosinophilic asthma

**DOI:** 10.1101/247171

**Authors:** Bart Hilvering, Timothy SC Hinks, Linda Stöger, Emanuele Marchi, Maryam Salimi, Rahul Shrimanker, Wei Liu, Wentao Chen, Jian Luo, Simei Go, Timothy Powell, Jennifer Cane, Samantha Thulborn, Ayako Kurioka, Tianqi Leng, Jamie Matthews, Clare Connolly, Catherine Borg, Mona Bafadhel, Christian B Willberg, Adaikalavan Ramasamy, Ratko Djukanović, Graham Ogg, Ian D Pavord, Paul Klenerman, Luzheng Xue

## Abstract

The functions and *in vivo* roles of type-2 CD8^+^ T cells in humans have not been well defined and this cell type has been largely overlooked in models of disease. We investigated this in the context of severe asthma with persistent airway eosinophilia - a phenotype associated with high exacerbation risk and responsiveness to type-2 cytokine-targeted therapies. In two independent cohorts we show that, in contrast to Th2 cells, type-2 cytokine-secreting CD8^+^CRTH2^+^ (Tc2) cells are enriched in blood and airways in severe eosinophilic asthma. Concentrations of prostaglandin D_2_ (PGD_2_) and cysteinyl leukotriene E_4_ (LTE_4_) are also increased in the airways of the same group of patients. *In vitro* PGD_2_ and LTE_4_ function synergistically to trigger Tc2 cell recruitment and activation in a TCR-independent manner. These lipids regulate diverse genes in Tc2 cells inducing type-2 cytokines and many other pro-inflammatory cytokines and chemokines which could contribute to eosinophilia. These findings are consistent with an important innate-like role for human Tc2 cells in severe eosinophilic asthma and suggest a potential target for therapeutic intervention in this and other diseases.

## Introduction

Type-2 cytokines (IL-4/5/9/13) orchestrate allergic inflammation, driving type-2 CD4^+^ T helper (Th2) cell differentiation, IgE production, mucus hypersecretion and airway hyperresponsiveness (AHR). Specifically, IL-5 activates and is chemotactic to eosinophils and prolongs their survival. Anti-type-2 cytokine therapies, notably mepolizumab, an anti-IL-5 antibody, are effective in severe eosinophilic asthma by reducing circulating eosinophils and asthma exacerbations^1–3^. The major sources of such type-2 cytokines are Th2, group 2 innate lymphoid cells (ILC2)^4^ and type-2 CD8^+^ T cells (Tc2). Of these, most attention has been paid to CD4^+^ T cells and more recently ILC2s, especially in human disease. Although, it has been known that type-2 CD8^+^ T cell populations exist, their overall functionality, transcriptional machinery and the mechanisms by which they are triggered have not been defined. This is important to address as recent data in other contexts have revealed previously overlooked functional diversity of human CD8^+^ T cells in inflammatory diseases^5^.

Eosinophilic asthma constitutes an important clinical phenotype, defined by increased airway eosinophils^6,7^ which release granule-derived basic proteins, lipid mediators, cytokines and chemokines, driving inflammation and exacerbations^8,9^. In some patients with severe asthma, airway eosinophils persist despite use of high-dose inhaled corticosteroids, suggesting relative steroid-insensitivity^10^. This phenotype is commonly associated with co-morbid rhinosinusitis, nasal polyposis and aspirin-induced bronchoconstriction^11^. Eosinophilic asthma is commonly considered as a Th2 disorder based on human data in mild asthma^12,13^ and animal models^14^. Recently ILC2s have been implicated in murine airway inflammation^15^, and increased ILC2s are reported in human asthma^16,17^. In contrast, although some data exists for overall involvement of CD8^+^ cells in asthma in both human^18,19^ and murine^20^ studies, the specific functional role of Tc2 cells remains largely unexplored, particularly in defined asthma phenotypes. Improved understanding of the pathogenic roles of Tc2 in this specific phenotype is important for therapeutic advances.

All type-2 cytokine-producing cells highly express chemoattractant receptor-homologous molecule expressed on Th2 cells (CRTH2), a receptor for prostaglandin D_2_ (PGD_2_)^4,21^. Through CRTH2, PGD_2_ elicits chemotaxis, type-2 cytokine production and suppresses apoptosis in Th2 and ILC2s^22–24^. The clinical efficacy of CRTH2 antagonists varies, being greatest in severe eosinophilic asthma^25,26^. We have previously shown synergistic enhancement of PGD_2_ with cysteinyl leukotrienes (cysLTs) in activating Th2 and ILC2s^27,28^. These lipid mediators and their receptors have not been studied in relation to CD8^+^ cells.

To investigate this, we first analysed type-2 CD8^+^ T cell frequencies and functional profiles in blood, bronchoalveolar lavage (BAL) and bronchial biopsies (BB) in well-defined patient cohorts, and further evaluated whether the airway environment is conducive to Tc2 activation via CRTH2 by measuring airway PGD_2_ and LTE_4_. We then defined the potent and broad activity of these lipids on Tc2 cells *in vitro* and investigate a mechanistic link between Tc2 cell activation and airway eosinophilia. Our observations provide compelling evidence of innate-like activation of Tc2 cells by pro-inflammatory lipids, a diverse range of functions of this cell population, and a potential role in severe eosinophilic asthma.

## Results

### Tc2 cells are enriched in eosinophilic asthma

CRTH2 is highly expressed on type-2 cytokine-producing human peripheral blood CD8^+^ T lymphocytes (described here as Tc2 cells) (Fig. 1a)^21^. We therefore first analysed human Tc2 cells using the phenotypic expression of CRTH2 on CD8^+^ T cells to define the Tc2 population in blood (Supplementary Fig. 1a). In a cohort of 56 participants from Oxford, UK, peripheral blood CD3^+^CD8^+^CRTH2^+^ Tc2 cells were substantially higher in patients with severe eosinophilic asthma (~6.24±5.18 % of CD8, n=26) than in severe non-eosinophilic asthma (~2.93±2.46 % of CD8, n=14, *p*<0.05) or health (~1.32±0.79 % of CD8, n=16, *p*<0.001) (Fig. 1b; Table 1). Findings were similar when blood Tc2 cells were expressed as a percentage of total leukocytes or in absolute numbers (Supplementary Fig. 1b,c; Supplementary Table 1). Conversely, the frequencies of CD3^+^CD4^+^CRTH2^+^ Th2 cells (Supplementary Fig. 1a) did not differ significantly (Fig. 1b; Supplementary Fig. 1b,c). Analysis of functional IL-5 and IL-13 producing CD8^+^ T cells *ex vivo* detected with PrimeFlow assays at mRNA level (Fig. 1c) and intracellular cytokine staining (ICS) at protein level (Fig. 1d; Supplementary Fig. 2) also supported Tc2 enrichment in severe eosinophilic asthma, although only small numbers of IL-5/IL-13 positive cells were detected by ICS in samples without stimulation (Supplementary Fig.2a). Furthermore, Tc2 cells were also found in the BAL and sputum in severe eosinophilic asthma (Supplementary Fig 3a). CD8+ T cells were detected abundantly in BB from the same group of patients (Supplementary Fig. 3b).

**Fig. 1.**
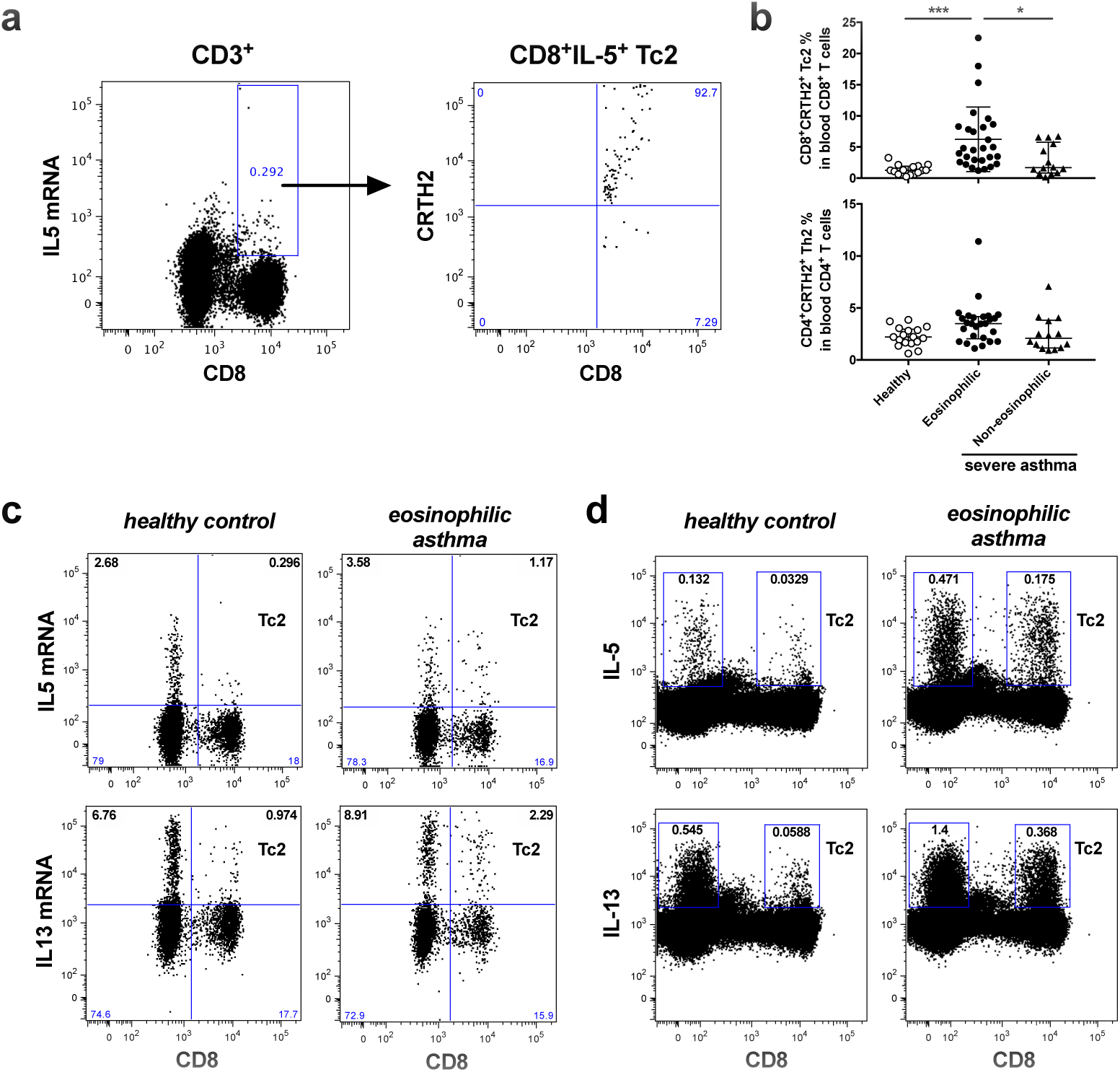
Tc2 cells are enriched in peripheral blood in severe eosinophilic asthma of the Oxford cohort. **a** IL-5-producing CD3^+^CD8^+^ Tc2 cells in human peripheral blood *ex-vivo* are CRTH2 positive **b**FrequenciesofCD3^+^CD8^+^CRTH2^+^Tc2and CD3^+^CD4^+^CRTH2^+^ Th2 cells in peripheral blood were compared between healthy controls and severe asthma patient groups (eosinophilic and non-eosinophilic) with flow cytometry. **c,d** IL-5- and IL-13-producing Tc2 cells in blood detected with PrimeFlow (**c**) or ICS after stimulation with 25 ng/ml PMA and 1µg/ml ionomycin (**d**) were compared between healthy control and severe eosinophilic asthma. \**p* < 0.05, \*\*\**p* < 0.001, (Data in **a**, **c** and **d** are representative of 3-5 independent experiments).

**Table 1.**
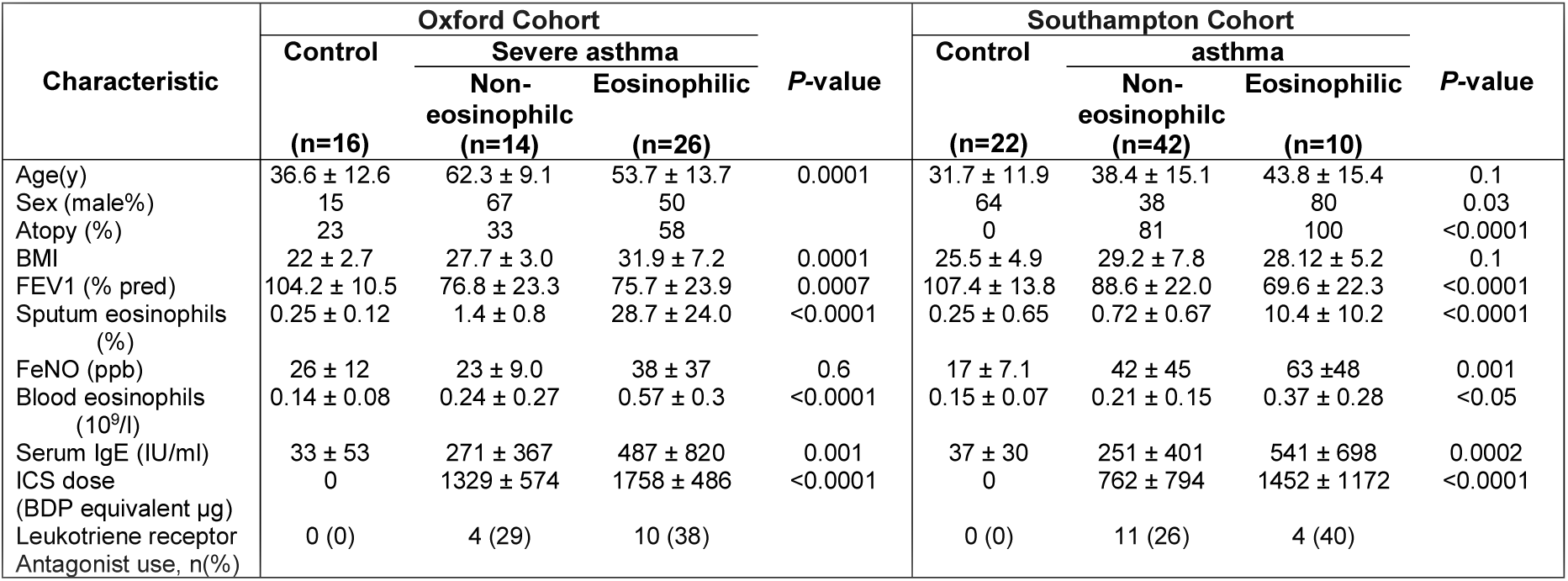
Study subjects (Mean ± SD)

We sought to confirm the validity of these findings in a second independent cohort of 74 participants from Southampton, UK (Table 1)^29^, using a different and complementary cell gating strategy, ICS (Supplementary Fig. 4a), which allows us to test whether the findings obtained using phenotypic staining were recapitulated using a functional assay. We found CD3^+^CD8^+^IL-13^+^ cells were increased in peripheral blood in asthma (~0.3% of CD8^+^ T cells, n=47) compared with health (~0.05%, n=19, *p*=0.04) (Supplementary Fig. 5a left panel). This increase correlated with asthma severity (Supplementary Fig. 5a middle-left panel, *p*=0.01), with the presence of nasal polyps (Supplementary Fig. 5a middle-right panel, *p*=0.008) and with previous smoking history (Supplementary Fig. 5a right panel, *p*=0.008), comorbidities associated with severe asthma. By contrast, frequencies of CD3^+^CD4^+^IL-13^+^ cells were significantly increased in mild (steroid-naïve) asthma (0.5%, n=14) compared with health (0.19% n=22, *p*<0.01), but not in steroid-treated moderate or severe asthma (Supplementary Fig. 5b). IL-4 expression in sputum T cells was associated positively with peripheral blood CD3^+^CD8^+^IL-13^+^ cell frequencies (*r*_*s*_=0.537, *p*=0.006), but negatively with peripheral blood CD3^+^CD4^+^IL-13^+^ cell frequencies (*r*_*s*_= -0.442, *p*=0.03) (Supplementary Fig. 6a).

As with peripheral blood (Fig. 1 and Supplementary Fig. 1b,c), CD3^+^CD8^+^IL-13^+^ cells (Supplementary Fig. 4b,c) were strikingly increased in BB and BAL in eosinophilic asthma (~2.05%, n=8 for BB; ~1.5%, n=9 for BAL amongst CD8^+^ T cells) compared with non-eosinophilic asthma (~0.1%, n=24 for BB; ~0.2%, n=26 for BAL) and health (~0.2%, n=13 for BB; ~0.2%, n=17 for BAL; *p*<0.005) in the Southampton cohort (Fig. 2). Again, CD3^+^CD4^+^IL-13^+^ cells in BB or BAL were not significantly increased in eosinophilic or non-eosinophilic forms. Amongst asthmatics, high frequencies of BB CD3^+^CD8^+^IL-13^+^ cells were associated with high bronchodilator reversibility (Supplementary Fig. 6b, *p*<0.007).

**Fig. 2.**
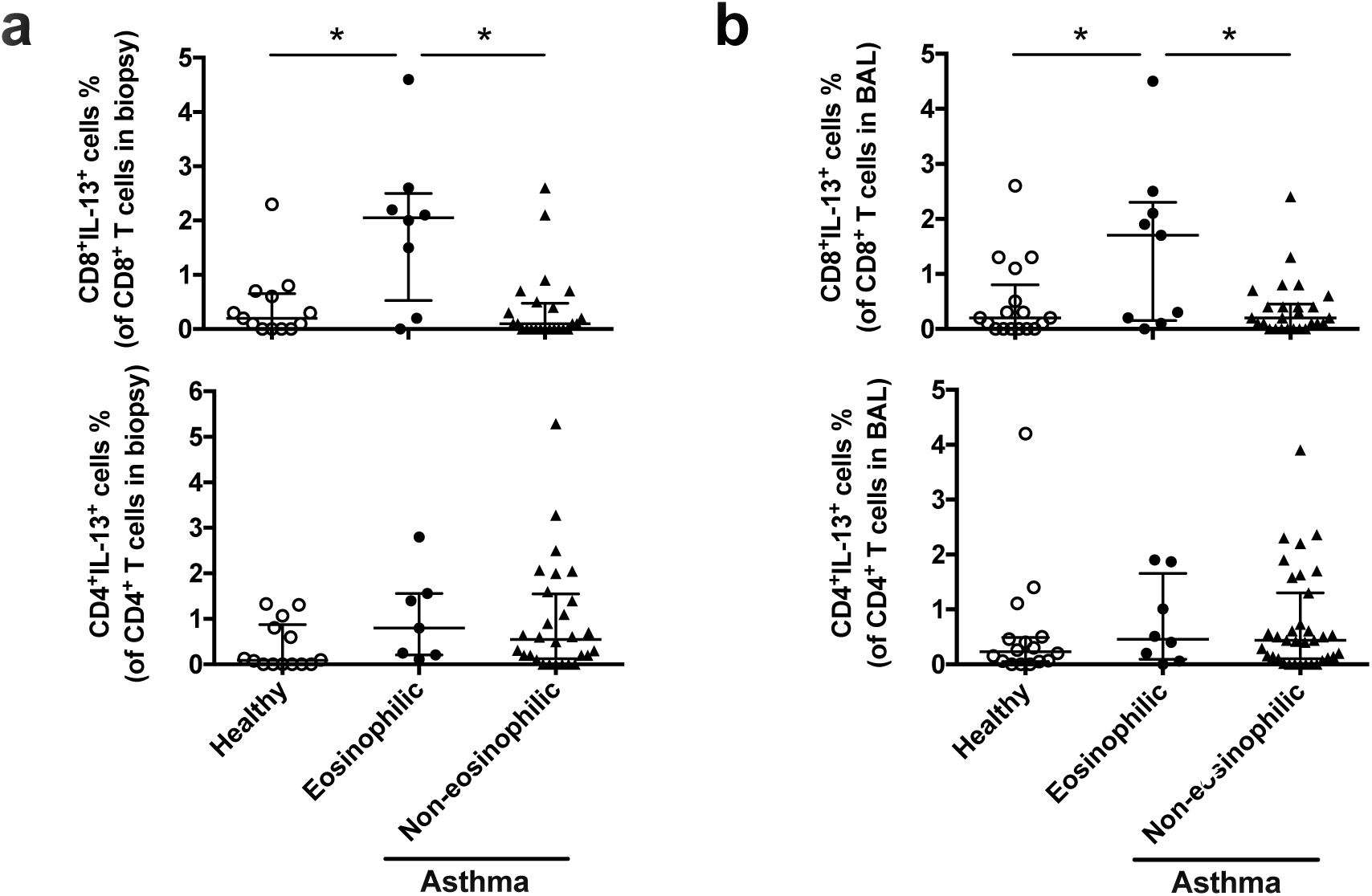
Tc2 cells are increased in lung in eosinophilic asthma of the Southampton cohortFrequencies of CD3^+^CD8^+^IL-13^+^ and CD3^+^CD4^+^IL-13^+^ T cells in BB (**a**) and BAL (**b**) were compared between healthy controls and asthma groups (eosinophilic and non-eosinophilic with ICS. \**p* < 0.05.

### Stimulatory eicosanoid mediators but not their receptors are enriched in eosinophilic asthma

Since Tc2 cells highly express CRTH2 together with CysLT_1_, a leukotriene receptor (Fig. 1a; Supplementary Fig. 7a), we compared their expression and levels of their ligands in the airways between asthma phenotypes (Fig. 3). The expression levels of CRTH2 and CysLT_1_ in individual Tc2 cells by flow cytometry were not significantly changed in the Oxford cohort (Fig. 3a). PGD_2_ assessed in sputum supernatants from asthma was significantly increased compared with health (2.76±0.54 ng/g, n=6, *p*<0.01) although no significant difference between eosinophilic (5.4±0.59 ng/g, n=11) and non-eosinophilic groups (7.2±1.46 ng/g, n=6, *p*=0.14) was detected (Fig. 3b). Pulmonary LTE_4_ was only significantly increased in patients with severe eosinophilic asthma (164.5±52 ng/g, n=10) but not in non-eosinophilic patients (1.84±0.65 ng/g, n=8), compared with health (0.34±0.15 ng/g, n=5, *p*=0.1) (Fig. 3b).

**Fig. 3.**
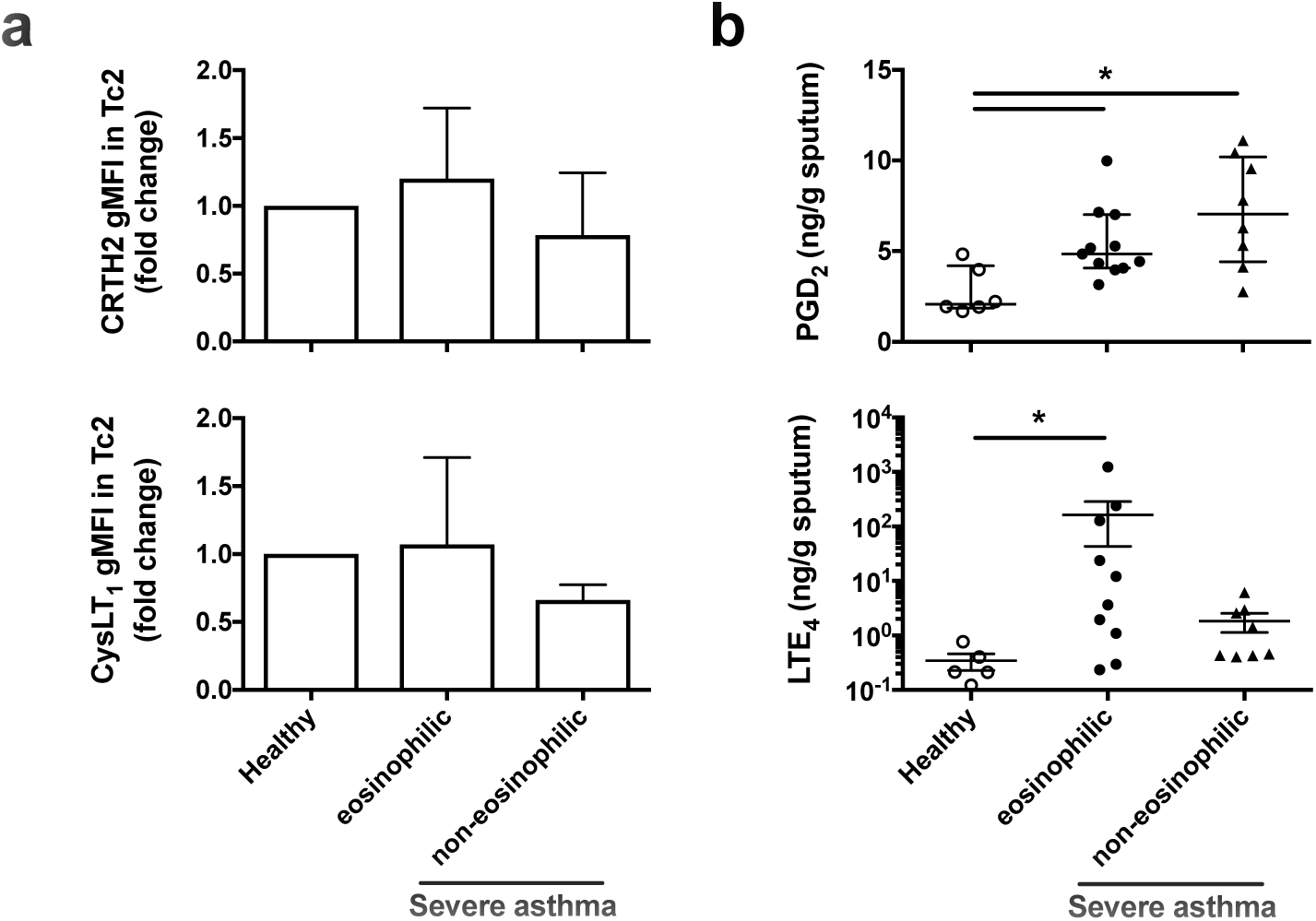
Eicosanoid mediators but not their receptors are increased in lung in severe eosinophilic asthma in the Oxford cohort. **a** Expressions of CRTH2 (upper panel) and CysLT_1_ (bottom panel) in individual Tc2 cells were paired compared between healthy controls and severe asthma groups with flow cytometry. **b** Levels of PGD_2_ (upper panel) and LTE_4_ (bottom panel) in sputum determined with ELISA were compared between healthy controls and severe asthma groups. \**p* < 0.05, n=5 for **a**.

### Tc2 cell migration induced by PGD_2_ and LTE_4_

To explore the potential pathogenic role of Tc2 cells and these lipid mediators, we isolated and cultured human Tc2 cells for further *in vitro* investigation (Supplementary Fig. 7b). These cells are CCR7^-^CD62L^-^ effectors (Supplementary Fig. 7c) and showed higher baseline type-2 gene expression signatures (*IL17RB, GPR44, CLECL1, IL9R, NAMPT, AF208111, HPGDS, P2RY14, RG4, IRS2* and *GATA3*) but lower type-1 (*IFNG, AIF1, LTA, TXK* and *IL18RAP*) and killer cell gene signatures (*KIR2DL1, KIR2DL4, KIR2DL5A, KLRF1, CD160* and *TYPOBP*) compared with other CD8^+^ cells (Supplementary Fig. 7d).

PGD_2_ and cysLTs are chemotactic agents for many types of immune cells^24,27^. To investigate airway Tc2 cell recruitment, we examined the effect of these lipids in chemotaxis assays. Both lipids caused cell migration in a typical bell-shaped dose-dependent manner, peaking at ~30 nM for PGD_2_ and ~10 nM for LTE_4_ (Fig. 4a). The maximum response induced by PGD_2_ was higher (2.4 fold) than that by LTE_4_. Cell migration was synergistically enhanced by combined stimulation (Fig. 4b). The contribution of PGD_2_ and LTE_4_ on the cell migration was blocked by the CRTH2 antagonist TM30089 and the CysLT_1_ antagonist montelukast.

**Fig. 4.**
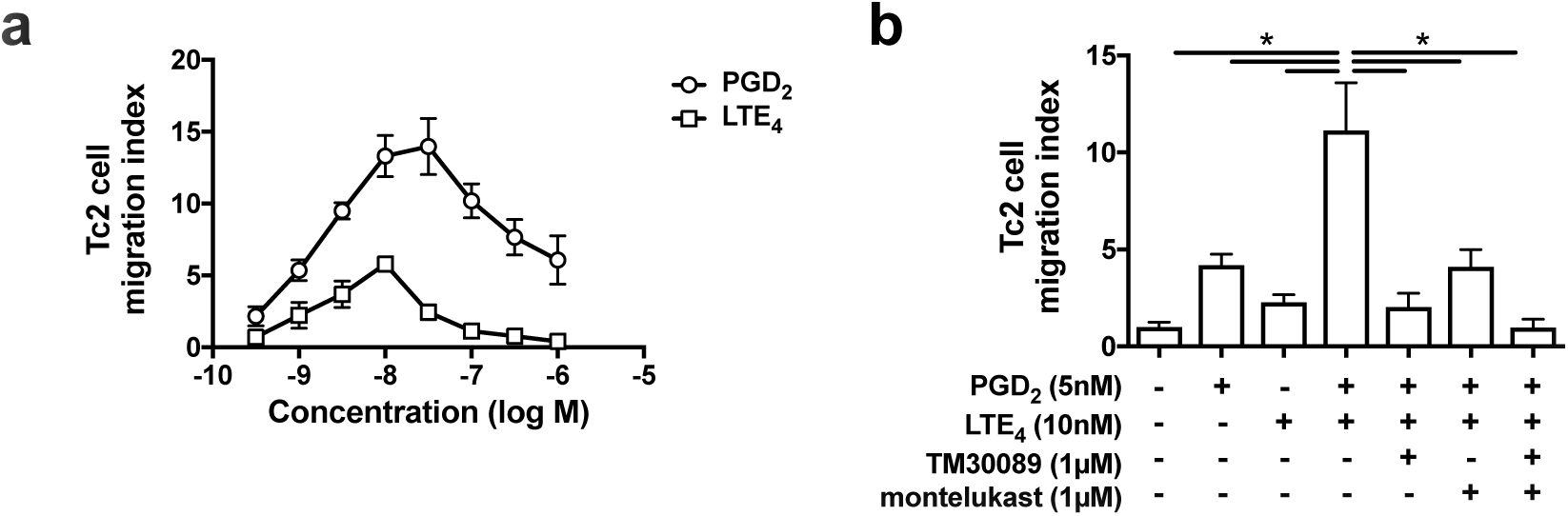
Tc2 cells (cultured) migrate in response to PGD_2_and LTE_4_. **a** Cell migration in responseto various concentrations of PGD_2_ and LTE_4_ in chemotaxis assays. **b** Cell migration in response to the combination of PGD_2_ and LTE_4_ in the absence or presence of TM30089 and montelukast. \**p* < 0.0001, (n=3).

### Enhancement of type-2 cytokine production in Tc2 cells by PGD_2_ and LTE_4_

We investigated the effects of PGD_2_ or LTE_4_ on type-2 cytokine production by Tc2 cells (Fig. 5). After treatment with PGD_2_ or LTE_4_, IL-5 and IL-13 production was elevated at both transcriptional (Fig. 5a) and translational (Fig. 5b) levels in a dose-dependent manner with EC_50_=17.3 nM (6.1 ng/ml) at mRNA or 17.9 nM (6.3 ng/ml) at protein on IL-5, and 21.1 nM (7.4 ng/ml) at mRNA or 16 nM (5.6 ng/ml) at protein on IL-13 for PGD_2_; and EC_50_=4.5 nM (2 ng/ml) at mRNA or 13.5 nM (5.9 ng/ml) at protein on IL-5, and 7.4 nM (3.3 ng/ml) at mRNA or 9 nM (4 ng/ml) at protein on IL-13 for LTE_4_, which were close to the concentrations of these lipids detected in patients’ sputa (Fig. 3b). Responses to PGD_2_ were significantly stronger than to LTE_4_. Compared with Th2 cells^22,30^, the effect of PGD_2_ and LTE_4_ on Tc2 is much more potent (Fig. 5c).

We then further examined type-2 cytokine production by Tc2 cells in response to the lipids alone or combination (Fig. 5d,e). Both PGD_2_ and LTE_4_ increased type-2 cytokine production. Their combination enhanced the response synergistically. Using PrimeFlow assays to analyse IL-5/IL-13 mRNA at individual cell level confirmed these data (Fig. 5f). IL-5/13 positive cells were increased from 2.88% (control) to 23.17%, 12.23% or 28.8% after treatment with PGD_2_, LTE_4_ or their combination respectively. Among these positive cells, some expressed IL-5 dominantly, some IL-13 dominantly, and only some produced IL-5/13 simultaneously, although >90% of these cells were capable of producing both following PMA/ionomycin stimulation (Supplementary Fig. 7e).

**Fig. 5.**
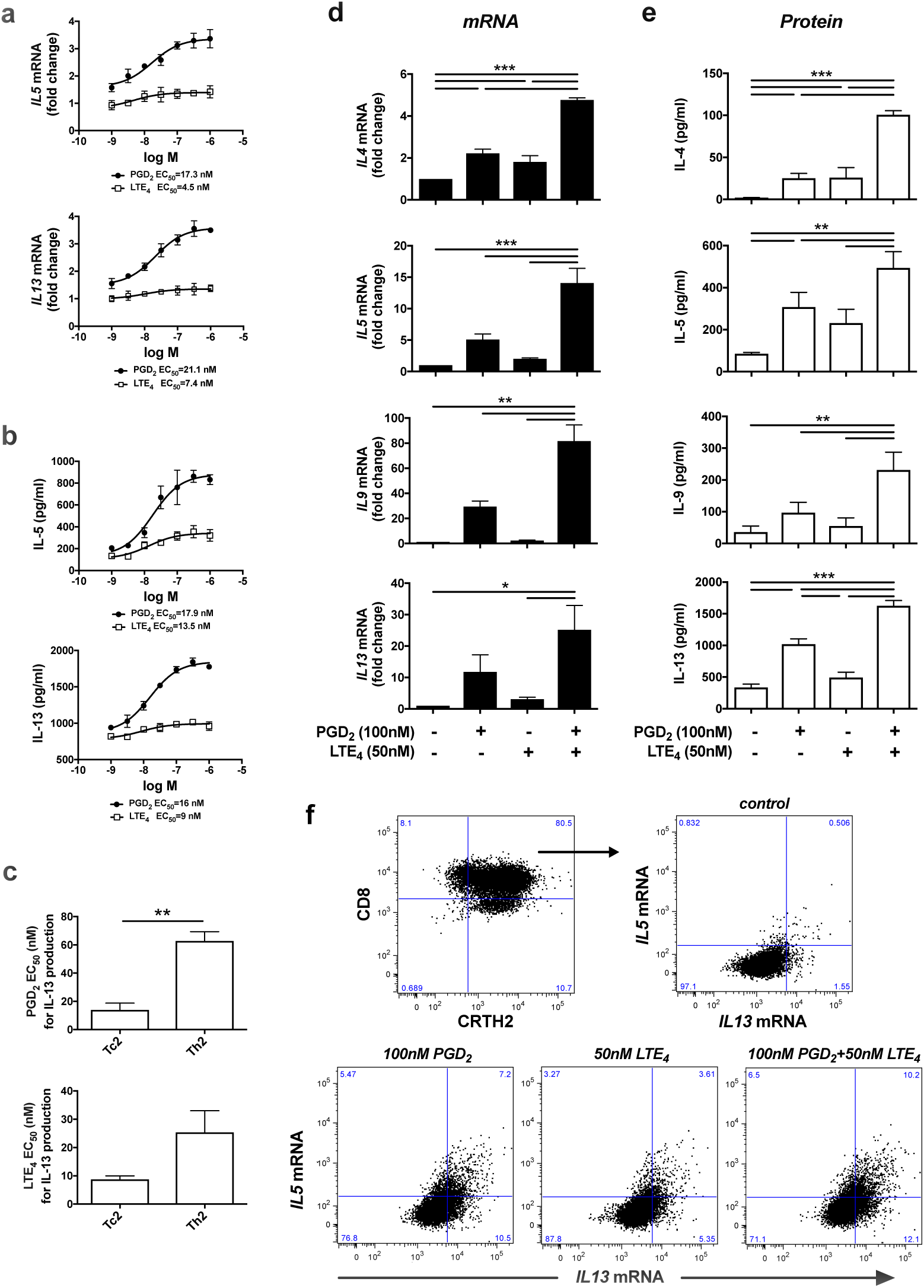
PGD_2_and LTE_4_promote type 2 cytokine production in cultured Tc2 cells. **a, b** mRNAlevels by qPRC in Tc2 cells (**a**) and protein levels by ELISA in the cell supernatants (**b**) for IL-5 and IL-13 after treatment with various concentration of PGD_2_ and LTE_4_. **c** EC_50_ of PGD_2_ or LTE_4_ for IL-13 production in Tc2 cells compared with that in Th2 cells. **d, e** mRNA levels in Tc2 cells measured by qPCR (**d**) and protein levels in the cell supernatants measured by Luminex (**e**) for type 2 cytokines after treatment with PGD_2_ and LTE_4_ alone or their combination. The mRNA levels in control samples were treated as 1-fold. **f** Increase of *IL5*-and *IL13*-mRNA positive Tc2 cells after treatment with PGD_2_ and LTE_4_ alone or their combination detected by using PrimeFlow RNA assay. \**p* < 0.05, \*\**p* < 0.005, \*\*\**p* < 0.001; data in **F** are representative of 3 independent experiments; (n=3 for **a**-**c** and **f**; n=6 for **d** and **e**).

### Effect of PGD_2_ and LTE_4_ on the gene expression profile of Tc2 cells

Using microarrays we investigated Tc2 transcriptional responses to PGD_2_ and LTE_4_. 1104, 3360, and 4593 gene transcriptions were significantly modulated (*P*<0.05) (including up/downregulation) by LTE_4_, PGD_2_, or their combination respectively (Supplementary Fig. 8a). The effect of PGD_2_ was much broader and stronger than that of LTE_4_, and the effect of the combination treatment was mainly contributed by PGD_2_ (Supplementary Fig. 8b).

We next focused on the genes encoding cytokines, chemokines, their receptors, and CD molecules (Fig. 6a; Supplementary Table 2). About 90 such genes were significantly modulated, mostly upregulated, and most obviously cytokines (*IL3*, *IL5*, *IL8*, *IL13*, *IL22*, *CSF2*, *TNF* and *XCL1*). Although a few of these were induced by LTE_4_ alone, most were driven by PGD_2_ alone or the combination. Some transcriptional changes were regulated only by the combination treatment (eg, *IL21*, *IL22*, *LASS1*, *CCL3*, *CCL4*, *IL1RL1*, *CD1E* and *CCL21*). Microarray data were largely confirmed by PCRarray on human common cytokines although some significant effects (*IL9* and *CSF1*) were detected only in PCRarray (Fig. 6b).

**Fig. 6.**
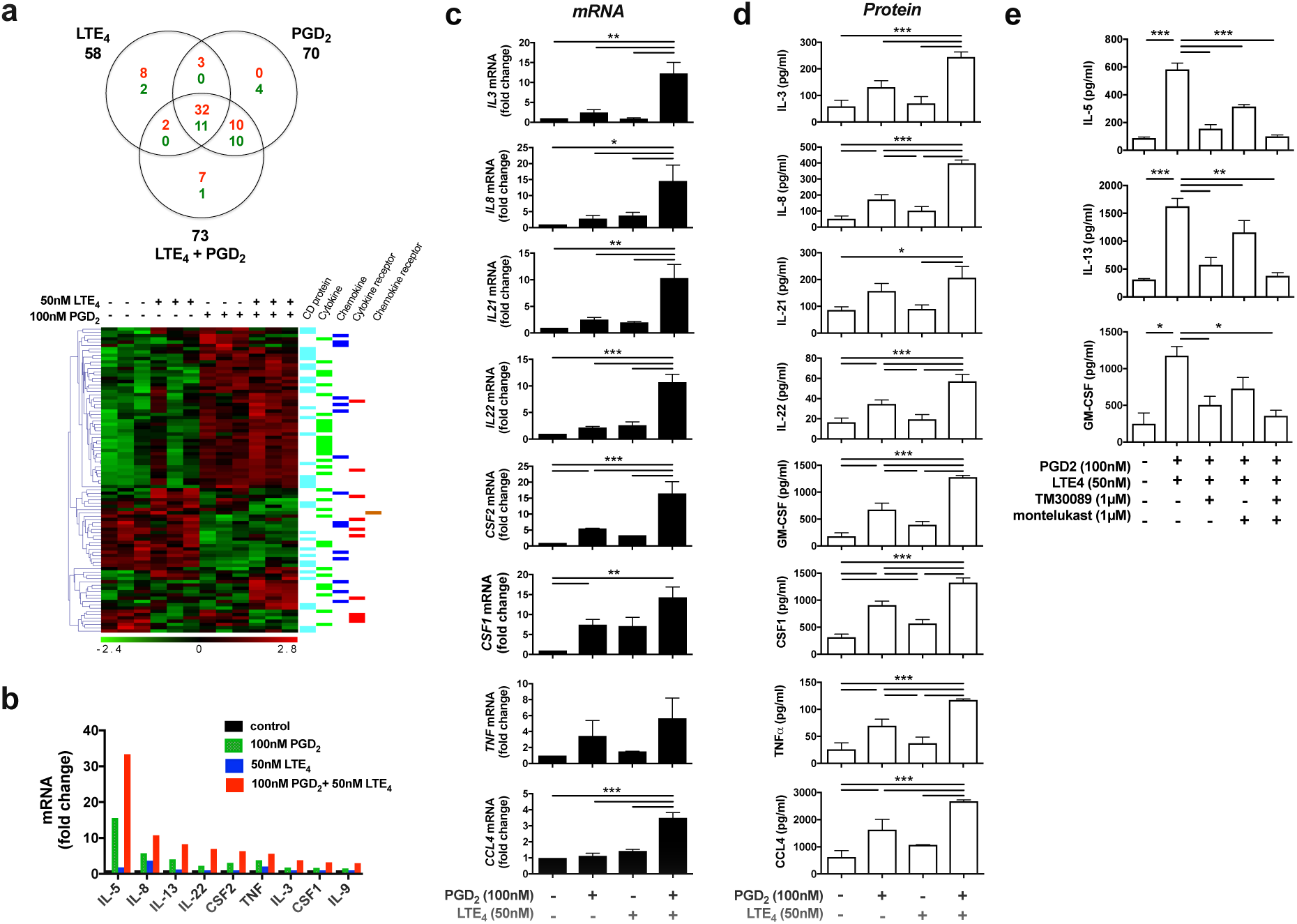
PGD_2_and LTE_4_modulated gene transcription and production of cytokines, chemokines,and surface receptors in cultured Tc2 cells. **a** Venn diagram and heat map showing significantly (*p* < 0.05) regulated genes for cytokines, chemokines and surface receptors (red for up-regulation and green for down-regulation) in Tc2 cells detected by microarray after treatments with PGD_2_, LTE_4_ or their combination. Three experimental replicates were prepared for each condition. **b** Up-regulated cytokine genes determined by using a PCRarray after the same treatments as those in (**a**). **c, d** The change of mRNA levels in cells measured with qPCR (**c**) and protein levels in the cell supernatants detected by using Luminex assays (**d**) for selected cytokines and chemokine after the same treatments as above. The mRNA levels in control samples were treated as 1-fold. **e** The cytokine production induced by PGD_2_ and LTE_4_ were inhibited by TM30089 and montelukast. \**p* < 0.05, \*\**p* < 0.005, \*\*\**p* < 0.001, (n=1 for **a**; n=3 for **b** and **c**; n=6 for **d**; n=4 for **e**).

For verification, we assayed selected cytokines by q-PCR (Fig. 6c) and Luminex (Fig. 6d). At mRNA level, most of genes (except *CSF1*) showed synergistic effects of PGD_2_ and LTE_4_. At protein level, effects of LTE_4_ were marginal in some cytokines (IL-3, IL-21 and IL-22), while effects of PGD_2_ were obvious in all the genes, particularly significant in IL-8, IL-22, GM-CSF, CSF1, TNFα and CCL4. Combination treatment either additively (CSF1) or synergistically (other genes) enhanced cytokine production. IL-2 enhanced the effects of the lipids on some cytokine (IL-5/8/13 and GM-CSF) production, particularly in PGD_2_ stimulation (Supplementary Fig. 9). The effects of PGD_2_ and LTE_4_ were inhibited by TM30089 and montelukast respectively (Fig. 6e).

Tc2 cells expressed cytotoxic proteins perforin and granzymes (GZMA, GZMB and GZMK) without stimulation (Supplementary Fig. 10). PGD_2_ and LTE_4_ had no significant effect, although activation of the cells by PMA/ionomycin downregulated the expression of the cytotoxic proteins.

### Effect of mast cell-derived PGD_2_ and LTE_4_ on the activation of Tc2 cells

To investigate the mechanism of Tc2 activation under more physiological conditions, we evaluated the impact of endogenously synthesized eicosanoids on Tc2 function. *In vitro* derived human mast cells were stimulated with IgE followed by crosslinking using an anti-IgE antibody (Fig. 7a). Only low levels of PGD_2_ (~0.1 ng/ml) and LTE_4_ (~6 ng/ml) were detected in mast cell supernatants before stimulation (supernatant UM). Supernatant from activated mast cells (supernatant SM) contained high levels of PGD_2_ (~11.5 ng/ml) and LTE_4_ (~86.8 ng/ml), which were similar to concentrations in sputum from asthmatic patients (Fig. 3b). Supernatant SM induced Tc2 cell migration (Fig. 7b) and marked cytokine (IL-5/13) production (Fig. 7c). Both effects were reduced by TM30089 and montelukast, and completely inhibited by their combination.

**Fig. 7.**
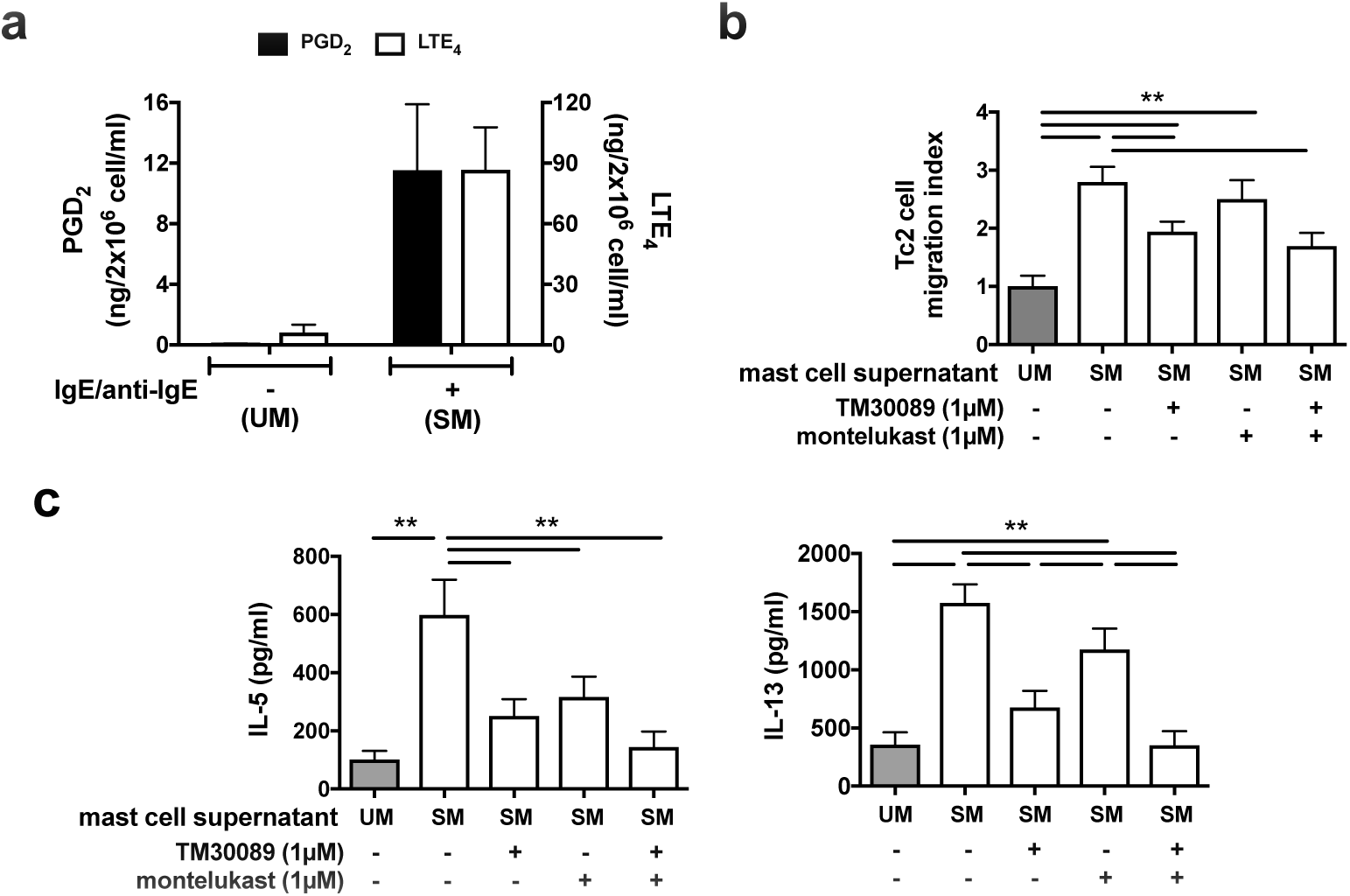
Tc2 cell activation in response to mast cell supernatant containing endogenous PGD_2_and LTE_4_ is mediated by receptors sensitive to TM30089 and montelukast respectively. **a** Levels of PGD_2_ (black bars) and LTE_4_ (white bars) were increased in supernatants from mast cells stimulated with IgE and anti-IgE antibodies (supernatant SM) compared with supernatants from cells without stimulation (supernatant UM). **b** More Tc2 cells migration to supernatant SM (white bars) than to supernatant UM (grey bar) in a chemotaxis assay was reduced by TM30089 and montelukast. **c** IL-5 and IL-13 productions in Tc2 cells were significantly increased in response to supernatant SM (white bars) compared with that to the supernatant UM (grey bars), and were inhibited by TM30089 and montelukast. \**p*<0.05, \*\**p* < 0.005, (n=3 for **a** and **b**; n=4 for **c**).

### The role of Tc2 in eosinophilia

Tc2 conditioned media were used to investigate the potential role of Tc2 cells in eosinophilia (Fig. 8). Anti-CD3/CD28 Tc2 stimulation increased IL-5 and GM-CSF secretion from ~600 and ~1300 pg/ml respectively in unstimulated supernatants (supernatant UT) to ~1290 and ~2900 pg/ml in stimulated cell supernatants (supernatant 3/28) (Fig. 8a).

**Fig. 8.**
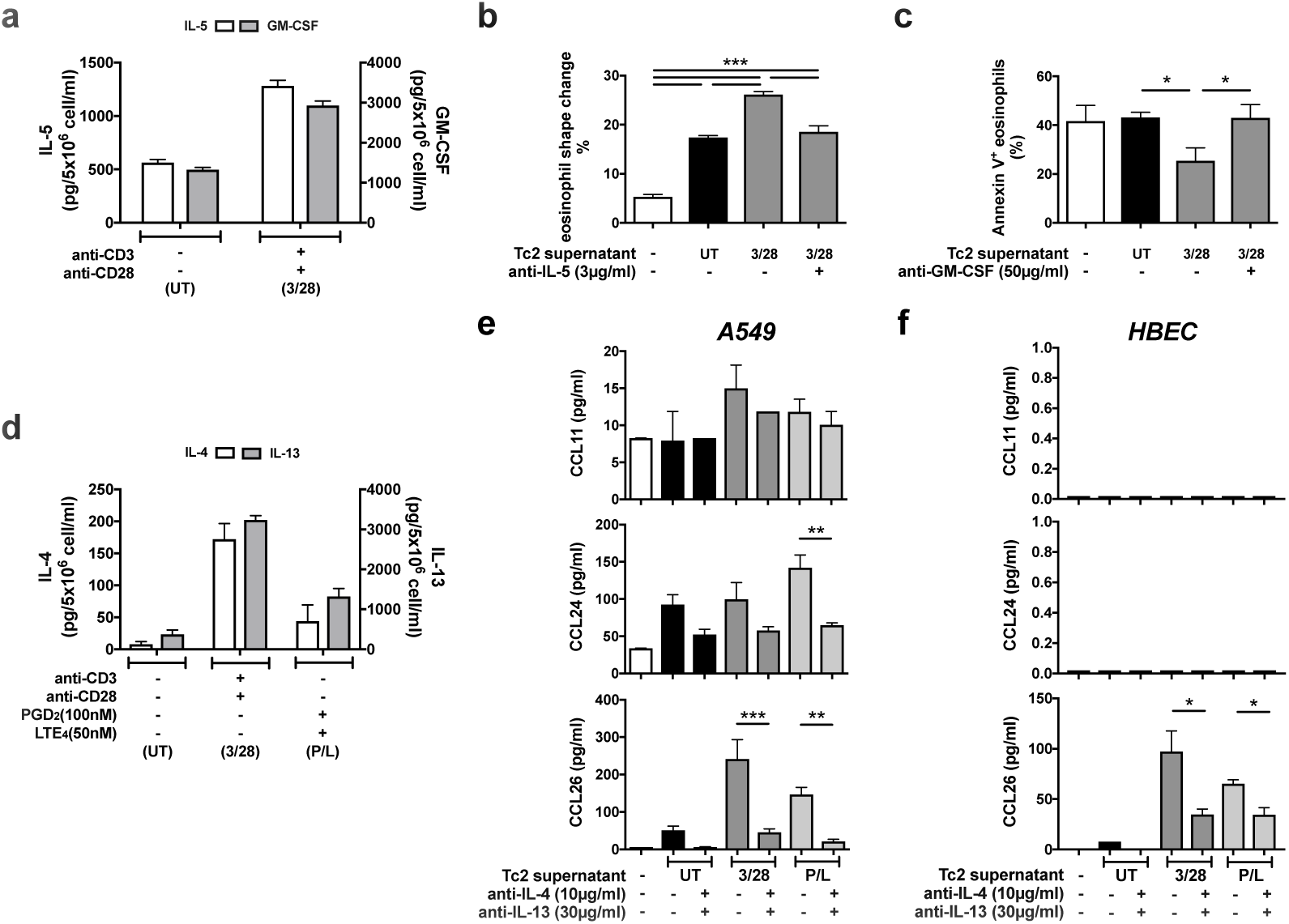
Cytokines released by human Tc2 cells promote eosinophil shape change and survival,and induce eosinophil chemokine production from airway epithelial cells. **a** Concentrations of IL-5 (white bars) and GM-CSF (grey bars) determined by ELISA were increased in the supernatants from Tc2 cells treated with anti-CD3/CD28 antibodies (supernatant 3/28) compared with the supernatants from the cells without treatment (supernatant UT). **b** Supernatant 3/28 induced more eosinophil shape change in fresh blood than supernatant UT, which was reversed by adding anti-IL-5 antibody. **c** Supernatant 3/28 reduced annexin V positive eosinophils induced by serum withdrawal in the culture, which was reversed by adding anti-GM-CSF antibody. **d** Concentrations of IL-4 (white bars) and IL-13 (grey bars) were increased in the supernatants of Tc2 cells treated with anti-CD3/CD28 antibodies (supernatant 3/28) or 100 nM PGD_2_/50 nM LTE_4_ (supernatant P/L) compared with untreated cells (supernatant UT). **e, f** Tc2 supernatants, particularly from treated cells, induced CCL11/24/26 (eotaxin-1/2/3) productions in A549 cells (**e**) or CCL26 in primary human bronchial epithelial cells (**f**) measured by Luminex, which were inhibited significantly by neutralizing antibodies against IL-4 and IL-13. \**p*<0.05, \*\**p* < 0.005, \*\*\**p* < 0.001, (n=3 for **a** and **f**; n=4 for **b-e**).

Recombinant human IL-5 (rhIL-5) induced eosinophil shape change, a biomarker of eosinophil migration, in fresh blood in a dose-dependent manner (Supplementary Fig. 11a), which was inhibited by anti-IL-5 neutralizing antibody. Supernatant 3/28 more potently induced eosinophil shape change *ex vivo* than supernatant UT (Fig. 8b). The reaction was inhibited partially by IL-5 antibody, implicating IL-5 and other eosinophil-active factors.

We examined the influence of GM-CSF released by Tc2 cells on eosinophil survival by measuring protection from serum starvation-induced apoptosis (Fig. 8c). The increase of Annexin V-positive (apoptotic) eosinophils after serum withdrawal was inhibited by rhGM-CSF, which was reversed by anti-GM-CSF neutralizing antibody (Supplementary Fig. 11b). Supernatant 3/28 provide similar protection, which was reduced by anti-GM-CSF antibody (Fig. 8c).

Eotaxins play a critical role in airway eosinophil recruitment in asthma^31^. Bronchial epithelial cells produce eotaxins, which is regulated by IL-4/13^32^. We investigated effects of IL-4/13 derived from Tc2 cells on eotaxin production in A549, a human alveolar epithelial cell line, and HBECs (Fig. 8d–f). Eotaxin generation, particularly eotaxin-2 (CCL24) and eotaxin-3 (CCL26), by A549 cells was induced by rhIL-4/13 (Supplementary Fig. 12a), and reversed by neutralizing antibodies against IL-4/13 (Supplementary Fig. 12b). Supernatant UT also up-regulated eotaxin production except eotaxin-1 (CCL11) in A549 (Fig. 8e). Supernatants from Tc2 cells activated by anti-CD3/CD28 antibodies (3/28) or PGD_2_/LTE_4_ (P/L) contain more IL-4/13 than supernatant UT (Fig. 8d), and showed higher capacities to induce eotaxins (Fig. 8e). These observations were confirmed in HBECs, although eotaxin-1/2 levels were not detected in these cells (Fig. 8f). Eotaxin induction by Tc2 media was significantly inhibited by anti-IL-4/13 neutralizing antibodies.

## Discussion

Tc2 cells are type-2 cytokine secreting CD8^+^ T lymphocytes that highly express CRTH2. We found enrichment of Tc2 cells in both peripheral blood and airways specifically in patients with severe asthma and persistent corticosteroid-insensitive eosinophilia in two independent cohorts. Importantly, the same conclusion was achieved using different methodologies, either surface CRTH2 expression or type-2 cytokine production as markers identifying Tc2 cells. Airway PGD_2_ and LTE_4_ were also increased in eosinophilic asthma. The interaction of Tc2 cells with these lipids induced strong activation of Tc2 cells, leading to cell migration and pro-inflammatory protein production, which in turn promoted eosinophil recruitment and survival directly or indirectly, suggesting an important role for Tc2 cells in the pathogenesis of eosinophilic asthma and their potential contribution to airway eosinophilia.

Research has long focussed on Th2 cells in asthma^13^ although blood Th2 cells are not associated with eosinophilia^33^, and bronchial Th2 cells are elevated in mild, but not severe asthma^29^. We found no significant differences in Th2 numbers between eosinophilic and non-eosinophilic asthma and healthy controls. Conversely Tc2 cells were significantly enriched, and associated with asthma severity, nasal polyposis, AHR and airway type-2 cytokines. This is consistent with reports in atopic asthma and non-atopic eosinophilic diseases^18,34^, of peripheral blood CD8^+^ T cell activation in severe asthma^35^ and observed associations between submucosal bronchial CD8^+^ cells and eosinophilia^36^. Our findings are also consistent with immunohistochemical findings of increased activation and IL-4 secretion by submucosal CD8^+^ T cells in fatal, virus-associated acute asthma^19^. Onset of severe eosinophilic asthma commonly follows a viral respiratory tract infection, which could promote CD8+ T cell activation including Tc2 cells leading to enhanced type-2 immunity in this phenotype. Furthermore, in murine models Th2 cell-derived IL-4 can reprogram virus specific CD8^+^ T cells to secrete type-2 cytokines^20,37^. Other murine data suggest preventing Tc2 cell differentiation reduces development of allergic airways disease^38,39^. Therefore, our findings strongly support the hypothesis that Tc2 cells, a previously unappreciated lymphocyte population, are key drivers contributing to type-2 immunity, particularly in severe eosinophilic asthma and should stimulate further investigation of these mechanisms.

Airway eosinophilia is associated with asthma exacerbations^40^ and airway remodelling^41^. In asthma, eosinophil granule proteins are cytotoxic and disrupt the protective pulmonary epithelial barrier triggering inflammation^42^. The importance of eosinophilic inflammation in asthma is best informed by response to therapies. Sputum eosinophilia is associated with responsiveness to corticosteroid therapy^43^. Other strategies such as anti-IgE, anti-IL-5 and CRTH2 antagonism aimed to normalize sputum eosinophils reduced exacerbation frequency and severity in clinical trials in asthma^2,3,25,26,40^. Our *in vitro* data have indicated that Tc2 activation can contribute to airway eosinophilia. Tc2-derived IL-5 and GM-CSF promote eosinophil migration and survival. Other Tc2 cytokines including IL-4/13 induce airway epithelial cells to produce eotaxins, the dominant chemokines for eosinophils in severe asthma, acting via CCR3^44^. Furthermore, in murine models, CD8^+^ T or virus-specific Tc2 cells mediate virus-induced lung eosinophilia and AHR^20,45^. All this evidence suggests an important role of Tc2 cells in eosinophil-mediated lung inflammation.

PGD_2_ and cysLTs are predominantly derived from mast cells in an IgE-dependent mechanism, which are abundant at sites of allergic responses, promoting airway inflammation and deterioration in lung function^46^. We observed increased airway PGD_2_ in eosinophilic and non-eosinophilic severe asthma, while LTE_4_ was enriched only in severe eosinophilic asthma, consistent with other reports^47^. Since many patients with severe eosinophilic asthma are non-atopic with low levels of IgE^6^, the key source of these lipids remains unclear. We previously demonstrated synergistic pro-inflammatory effects of PGD_2_ and LTE_4_ in human Th2 and ILC2s, which inhibited by TM30089 and montelukast^27,28,30^. This also occurs in Tc2 cells, indicating that these CD8 cells can be activated by lipid mediators in an antigen-independent manner. Of relevance, we observed higher potency of PGD_2_ and LTE_4_ on Tc2 than on Th2 cells. Synergistic effects of PGD_2_ and LTE_4_ in Tc2 were replicated using endogenous lipid mediators from activated mast cells. These findings provide a potential mechanistic insight into the clinical efficacy of CRTH2 antagonism in severe eosinophilic asthma^26^.

Transcriptional analysis suggested a broad range of effects of these lipids beyond type-2 cytokine induction in Tc2 cells, with >2000 upregulated transcripts including genes for bioactive cytokines and chemokines for eosinophils: IL-3, IL-5 and GM-CSF promote eosinophil differentiation and migration^48^; TNFα delays eosinophil apoptosis^49^; overexpression of IL-25 resulted in eosinophilia in murine models^50^; and CCL3, CCL4 and CCL7 are chemokines for eosinophils^51^. One notable finding was the impact of these lipids on IL-5 expression by Tc2 cells. The EC_50_ for PGD_2_ induced IL-5 production was 17.9 nM, compared to 63 nM for Th2 cells^22^. Thus Tc2 cells stimulated in this way generate a strong and early IL-5 response, which could be relevant to the clear clinical benefit of anti-IL-5 in severe eosinophilic asthma^3,40,52^.

In conclusion, Tc2 represent an important cell type with innate-like characteristics and substantial pro-inflammatory potential. *In vivo* there is significant enrichment of Tc2 cells in both peripheral blood and airways in eosinophilic asthma, particularly in severe disease, associated with increases in airway PGD_2_ and LTE_4_. These lipid mediators function potently and synergistically to recruit and activate Tc2 cells to produce type-2 and multiple other pro-inflammatory cytokines and chemokines, which sufficient to contribute to airway eosinophilia. Taken together these data suggest that Tc2 cells constitute potentially important therapeutic targets in severe eosinophilic asthma.

## Methods

### Reagents

PGD_2_ and LTE_4_ were purchased from Enzo Life Science; TM30089 (CAY10471) and montelukast were supplied by Cayman Chemical; rhIL-5, anti-CD3, anti-CD28 antibodies, human CD8^+^ T cell isolation kit and anti-human CD294 or CD16 MicroBeads were from Miltenyi Biotec Ltd; BD FACS Lysing Solution was supplied by BD Biosciences; X-VIVO 15 medium and Bronchial Epithelial Cell Growth Medium (BEGM) were purchased from Lonza; AIM V medium was from Invitrogen; Ficoll-PaqueTM Plus was supplied from GE Healthcare; RNeasy Mini kit and Omniscript reverse transcription kit were supplied from Qiagen; Real time quantitative PCR (qPCR) Master Mix and probes were from Roche; Primers were synthesized by Eurofins MWG Operon; rhSCF, rhIL-6 and anti-IL-4/5/13 neutralizing antibodies were purchased from Bio-techne; goat anti-human CD3 was from Santa Cruz; rabbit anti-human CD8 was from Abcam; peroxidase polymer-conjugated anti-rabbit and peroxidase polymer-conjugated anti-goat antibodies were from Vector Laboratories; Fluorescein-tyramide and Cy5-tyramide were from PerkinElmer; human myeloma IgE was from Calbiochem; Anti-human GM-CSF antibody, Annexin V-APC and Human GM-CSF ELISA kit were obtained from BioLegend; IC fixation buffer, Permeabilisation Buffer, human IL-4, IL-5 and IL-13 ELISA kit were from eBioscience; and rhIL-2, rhIL-4 and rhGM-CSF were from PeproTech and other chemicals were from Sigma-Aldrich.

### Human clinical samples

For the Oxford cohort, patients meeting the ATS/ERS definition of severe asthma^10^ with a sputum eosinophil count of >3% (eosinophilic, n=26) or <3% (non-eosinophilic, n=14), and 16 healthy control subjects were recruited from Churchill Hospital, Oxford (Table 1). For the Southampton cohort, 22 healthy participants, 10 with eosinophilic asthma (sputum eosinophil count >3%), and 42 non-eosinophilic asthma (Table 1) were enrolled from NIHR Southampton Respiratory Biomedical Research Unit and outpatient clinics at University Hospital Southampton^29^.

Peripheral blood was collected and used directly for flow cytometry. Sputum was induced with nebulized saline solution (3-5%) after pre-treatment with salbutamol. Selected sputum plugs were dispersed with 0.2% DTT, filtered for cells for flow cytometry analysis or microarray, and supernatants for ELISA analysis. BB and BAL were collected under bronchoscopy^29^. Biopsies were dispersed with collagenase for 1 h, and BALs were treated with 0.1% DTT and filtered for flow cytometry. For microarray, CD3^+^ sputum cells were FACS sorted using a FACS Aria II^TM^ cell sorter (BD Biosciences).

### Human CD8^+^CRTH2^+^ Tc2 cell preparation and treatment

Human Tc2 cells were isolated from CD Leucocyte Cones (National Blood Service, UK). Briefly, CD8^+^ cells were isolated from PBMC using MACS CD8^+^ T cell isolation kit, followed by CRTH2-positive selection using anti-human CD294 MicroBeads, and further amplified in X-VIVO 15 medium containing 10% human serum and 50 U/ml rhIL-2.

For gene or protein analysis, Tc2 cells were treated with conditions as indicated in the results for 4 h. Cell supernatants were collected for ELISA or Luminex assays, and cell pallets for qPCR, RNAarray or microarray.

For Tc2-conditioned supernatants, cells were treated with immobilized anti-CD3/anti-CD28 antibodies or PGD_2_ (100 nM)/LTE_4_ (50 nM) for 4 h, and then supernatants were harvested for ELISA, or used as Tc2-conditioned media for the treatment of fresh blood, eosinophils, A549 cells or primary human bronchial epithelial cells (HBECs).

### Human Mast cell culture and treatment

Human mast cells were cultured and treated as described previously^30^. Briefly, CD34^+^ progenitor cells were isolated from human cord blood (National Blood Service, Oxford, UK) by using a human CD34 MicroBead kit. The cells were cultured with Iscove’s modified Dulbecco’s medium containing 10% human serum, 0.55 µM 2-ME, penicillin/streptomycin, human recombinant stem cell factor (100 ng/ml) and human rIL-6 (50 ng/ml) for 14–15 weeks. Half the culture medium was replaced twice weekly with fresh medium containing the same concentration of cytokines. The cells were pre-treated with 5 mg/ml purified human myeloma IgE for 4 days, washed and then sensitized passively with fresh IgE (5 mg/ml) for 2 h. After washing, the cells were incubated with medium or challenged with goat anti-human IgE (1 µg/ml) for 1 h. The supernatants of the cells were collected and measured for PGD_2_ and LTE_4_, and used as mast cell supernatants for the treatment of Tc2 cells.

### Human bronchial epithelial cell culture and treatment

A549 cells (ATCC) were cultured in RPMI-1640 medium with 10% FCS. Primary HBECs were obtained from bronchial brushings collected under bronchoscopy, and cultured with Bronchial Epithelial Cell Growth Medium containing 50 µg/ml gentamicin and 1×penicillin/streptomycin. Both cell types were treated with rhIL-4 and rhIL-13 in RPMI 1640 (5% FCS) or Tc2 conditioned media in the presence or absence of anti-IL-4 and IL-13 neutralizing antibodies as indicated in the results for 16 h. The supernatants of the cells were harvested for Luminex assay for eotaxins.

### PrimeFlow RNA assay

The levels of transcription for IL-5 and IL-13 in individual Tc2 cells in whole blood or Tc2 cultures were analysed with a PrimeFlow RNA Assay kit (eBioscience) according to the manufacturer’s instructions. Briefly, fresh blood or purified Tc2 cells were treated with conditions indicated for 4 h, and then were stained with the antibodies (Table E3) and viability dye followed by fixation and permeabilisation. Then the cells were hybridised with RNA probes for IL-5 and IL-13. The signals were amplified and labelled with fluorescent probes. The results were analysed with a BD LSRFortessa flow cytometer (BD Biosciences).

### Microarrays

For CD3^+^ cells sorted from sputa, RNAs were isolated using an Absolutely RNA Nanoprep Kit (Agilent), and microarrays were performed using Affymetrix HT HG-U133+ PM GeneChips by Janssen Research & Development (Springhouse, Pennsylvania). For cultured Tc2 and CRTH2^-^CD8^+^ T cells, RNAs were extracted with an RNeasy Mini kit, and microarrays were conducted using an Illumina HumanHT-12v4 Expression Beadchip at the Transcriptomics Core Facility, The Jenner Institute, University of Oxford. Pre-processing data analysis was performed using R language (www.R-project.org) and Bioconductor packages (www.bioconductor.org/). Genes significant at a *p*<0.05 were selected by using Limma Bioconductor package^53^. Heat maps and gene hierarchal clustering were generated by using *tmev* microarray software suite.

### ELISA

The levels of IL-4, IL-5, IL-13, and GM-CSF in the Tc2 supernatants were assayed with ELISA kits, and the concentrations of PGD_2_ and LTE_4_ in the mast cell supernatants were measured with a PGD_2_–MOX enzyme immunoassay kit and LTE_4_ enzyme immunoassay kit (Cayman Chemicals) respectively according to the manufacturer’s instructions. The results were measured in an EnVision Multilable Reader (PerkinElmer).

### Luminex assays

Multiple cytokine concentrations in the supernatants of Tc2 cultures or multiple eotaxin concentrations in the supernatants of A549 cells and HBECs after various treatments as indicated were measured using a Luminex Screening Assay kit (Bio-techne) as per the manufacturer’s instructions. Results were obtained with a Bio-Plex 200 System (Bio-Rad).

### PCRarray

The levels of mRNA for cytokines in the RNA samples from the Tc2 cells after treatments were assessed with PCRarray by using an RT^2^ Profiler PCRarray Human Common Cytokines PCR Array kit (Qiagen) in a LightCycler 480 Real-Time PCR System (Roche).

### Quantitative RT-PCR (qPCR)

qPCR was conducted as described previously^30^. Briefly, total RNA of the cells was extracted using an RNeasy Mini kit. cDNA of the samples was prepared from the same starting amount of RNA using a Omniscript RT kit. qPCR was conducted using Master Mix and Probe in a LightCycler 480 Real-Time PCR System (Roche). GAPDH was used as control gene. Primers and probes (Roche) used are listed in Table E4.

### Flow cytometry

For blood samples, fresh blood was labelled using antibodies (Table E3) followed by red blood cell lysis with a BD FACS Lysing Solution. For cells from blood, sputum, BAL or BB, the cells were stained with antibody cocktail and live/dead dye (Table E3). The protocols have been optimised with intraclass correlation coefficient (ICC) = 0.86 for Tc2, 0.71 for Th2 and 0.96 for ILC2, and most of the experiments were paired between asthmatic patients and healthy controls. For intracellular cytokine staining in the Oxford cohort, PBMCs from fresh blood were treated without or with 25 ng/ml PMA and 1 µg/ml ionomycin in the presence of 5 µg/ml brefeldin A for 6 h. In the Southampton cohort, the cells were rested overnight in AIM-V medium and stimulated with 25 ng/ml PMA and 500 ng/ml ionomycin in the presence of 2 µM monensin for 5 h. After surface marker staining, the cells were fixed with IC fixation buffer (Oxford) or 1% formaldehyde (Southampton) and then treated with a Permeabilisation Buffer followed by incubation with anti-IL-5 and IL-13 antibodies. Fluorescence-minus-one (CRTH2, IL-5 and IL-13) controls (Oxford) or unstimulated controls (Southampton) were included in each experiment. The samples were analysed with a BD LSRFortessa flow cytometer at Oxford or a BD FACS Aria II cell sorter at Southampton.

### Chemotaxis assays

Tc2 cells were resuspended with RPMI 1640 media; 25 mL of cell suspension and 29-mL test compounds as indicated in the results or mast cell supernatants were applied to upper and lower chambers, respectively, in a 5-µm pore sized 96-well ChemoTx plate (Neuro Probe). After incubation (37°C for 60 minutes), the migrated cells in the lower chambers were collected and mixed with a Cell Titer-Glo Luminescent Cell Viability Assay kit (Promega) and quantified by using an EnVision Multilable Reader.

### Immunohistochemistry

Paraffin-embedded sections of BB were prepared by Oxford Centre for Histopathology Research. After deparaffin and rehydation, the sections were boiled in a Target Retrieval Solution (Dako), followed by incubation with peroxidase blocking reagent (Bio-Rad) and normal horse serum. The sections were then labelled with primary (goat anti-human CD3 and rabbit anti-human CD8) and secondary (peroxidase polymer-conjugated anti-rabbit followed by Cy5-tyramide) antibodies. After treatment with peroxidase blocking reagent again, the sections were further incubated with another secondary antibody (peroxidase polymer-conjugated anti-goat followed by Fluorescein-tyramide) and then 5 µg/ml DAPI solution. Images were acquired on an Olympus FV1200 inverted confocal microscope, and processed with ImageJ.

### Eosinophil shape change assay

Fresh human blood was incubated with an equal volume of Tc2 conditioned media in the presence or absence of anti-IL-5 neutralizing antibody or other reagents as indicated for 1 h. The samples were fixed with a Cytofix Fixation Buffer (BD Biosciences) followed by red blood cell lysis by using RBC Lysis Solution (Gentra Systems). Eosinophils were gated from granulocytes according to their autofluorescence during the analysis in a BD LSRFortessa flow cytometer. The eosinophil shape change was determined by the position shifting of the cells in forward scatter.

### Eosinophil apoptosis assay

Eosinophils were prepared from human blood. Erythrocyte/granulocyte pellet was collected after Ficoll-Hypaque gradient, and then incubated with 3% dextran saline solution for sedimentation. Granulocytes were harvested from the supernatant of the sedimentation and further purified by lysis of the remaining erythrocytes with 0.6 M KCl hypotonic water, and followed by labelling with anti-CD16 microbeads. Unlabelled eosinophils were negatively selected and then resuspended in RPMI 1640 medium. After treatments with diluted Tc2-conditioned media in the presence or absence of anti-GM-CSF neutralizing antibody or other reagents as indicated for 12 h, the eosinophils were labelled with Annexin V and then analysed with a BD LSRFortessa flow cytometer.

### Statistics

Data for clinical samples in Figures 1, 2, 3, and Supplementary Fig. 1, 4, 5 are presented as medians with interquartile range (IQR) and other data are presented as means with SEM. Data were analysed using one-way ANOVA followed by the Tukey’s test or Student’s t-tests. Groups ranked according to disease severity were tested for linear trends using Jonckheere-Terpstra tests. Values of *p*<0.05 were considered statistically significant.

### Study approval

The studies were approved by Leicestershire, Nottinghamshire Rutland Ethics Committee, UK (08/H0406/189) and the Southampton and South West Hampshire Research Ethics Committee B, UK (10/H0504/2).

## Acknowledgements

This work was supported by grants from European Respiratory Society (BH), Wellcome Trust (TSCH-088365/z/09/z, 104553/Z/14/Z, PK-WT109965MA), the Academy of Medical Sciences (TSCH), MRC (GO), NIHR Biomedical Research Centre Programme, Oxford (GO, IDP, PK, LX), NIHR Academic Clinical Fellowship (TSCH), NIHR Senior Fellowship (PK), Oxford Martin School (PK) and British Medical Association (The James Trust 2011) (GO, PK, LX).

## Author Contributions

BH, TSCH, LS, EM performed the experiments and collected, analysed, and discussed the data; MS, RS, WL, WC, JL, SG, JC, TP, ST, AK, TL, JM, CC, CB, MB, CW were involved in part of the experiments, patient recruitment and patient sample collection; AR conducted microarray; TSCH, RD, GO designed the experiments, and were involved in the manuscript writing; IP, PK, LX managed and designed the study, conceived the experiments and wrote the manuscript, which was critically reviewed and approved by all authors.

## REFERENCES

1. Hilvering, B., Xue, L., & Pavord, I. D. Evidence for the efficacy and safety of anti-interleukin-5 treatment in the management of refractory eosinophilic asthma. Ther Adv Respir Dis. 9, 135–145 (2015).

2. Haldar, P. et al. Mepolizumab and exacerbations of refractory eosinophilic asthma. N Engl J Med. 360, 973–984 (2009).

3. Ortega, H. G. et al. Mepolizumab treatment in patients with severe eosinophilic asthma. N Engl J Med. 371, 1198–1207 (2014).

4. Mjösberg, J. M. et al. Human IL-25- and IL-33-responsive type 2 innate lymphoid cells are defined by expression of CRTH2 and CD161. Nat Immunol. 12, 1055–1062 (2011).

5. Billerbeck, E. et al. Analysis of CD161 expression on human CD8+ T cells defines a distinct functional subset with tissue-homing properties. Proc Natl Acad Sci USA. 107, 3006–3011 (2010).

6. Haldar, P. et al. Cluster analysis and clinical asthma phenotypes. Am J Respir Crit Care Med. 178, 218–224 (2008).

7. Nair, P. What is an “eosinophilic phenotype” of asthma? J Allergy Clin Immunol. 132, 81–83 (2013).

8. Nissim Ben Efraim, A. H..& Levi-Schaffer, F. Tissue remodeling and angiogenesis in asthma: the role of the eosinophil. Ther Adv Respir Dis. 2, 163–171 (2008).

9. Bousquet, J. et al. Eosinophilic inflammation in asthma. N Engl J Med. 323, 1033–1039 (1990).

10. Chung, K.F. et al. International ERS/ATS guidelines on definition, evaluation and treatment of severe asthma. Eur Respir J. 43, 343–373 (2014).

11. Mullol, J..& Picado, C. Rhinosinusitis and nasal polyps in aspirin-exacerbated respiratory disease. Immunol Allergy Clin North Am. 33, 163–176 (2013).

12. Lambrecht, B. N..& Hammad, H. The immunology of asthma. Nat Immunol. 16, 45–56 (2015).

13. Robinson, D. S. et al. Predominant TH2-like bronchoalveolar T-lymphocyte population in atopic asthma. N Engl J Med. 326, 298–304 (1992).

14. Doherty, T. A., Soroosh, P., Broide, D. H. & Croft, M. CD4+ cells are required for chronic eosinophilic lung inflammation but not airway remodeling. Am J Physiol Lung Cell Mol Physiol. 296, L229–235 (2009).

15. Halim, T. Y., Krauss, R. H., Sun, A. C. & Takei, F. Lung natural helper cells are a critical source of Th2 cell-type cytokines in protease allergen-induced airway inflammation. Immunity 36, 451–63 (2012).

16. Bartemes, K. R., Kephart, G. M., Fox, S. J. & Kita, H. Enhanced innate type 2 immune response in peripheral blood from patients with asthma. J Allergy Clin Immunol. 134, 671–678 e4 (2014).

17. Nagakumar, P. et al. Type 2 innate lymphoid cells in induced sputum from children with severe asthma. J Allergy Clin Immunol. 137, 624–626 e6 (2016).

18. Cho, S. H., Stanciu, L. A., Holgate, S. T. & Johnston, S. L. Increased interleukin-4, interleukin-5, and interferon-gamma in airway CD4+ and CD8+ T cells in atopic asthma. Am J Respir Crit Care Med. 171, 224–230 (2005).

19. O'Sullivan, S. et al. Activated, cytotoxic CD8(+) T lymphocytes contribute to the pathology of asthma death. Am J Respir Crit Care Med. 164, 560–564 (2001).

20. Coyle, A. J. et al. Virus-specific CD8+ cells can switch to interleukin 5 production and induce airway eosinophilia. J Exp Med. 181, 1229–1233 (1995).

21. Cosmi, L. et al. CRTH2 is the most reliable marker for the detection of circulating human type 2 Th and type 2 T cytotoxic cells in health and disease. Eur J Immunol. 30, 2972–2979 (2000).

22. Xue, L. et al. Prostaglandin D2 causes preferential induction of proinflammatory Th2 cytokine production through an action on chemoattractant receptor-like molecule expressed on Th2 cells. J Immunol. 175, 6531–6536 (2005).

23. Xue, L., Barrow, A. & Pettipher, R. Novel function of CRTH2 in preventing apoptosis of human Th2 cells through activation of the phosphatidylinositol 3-kinase pathway. J Immunol. 182, 7580–7586 (2009).

24. Xue, L. et al. Prostaglandin D2 activates group 2 innate lymphoid cells through chemoattractant receptor-homologous molecule expressed on TH2 cells. J Allergy Clin Immunol. 133, 1184–1194 (2014).

25. Pettipher, R. et al. Heightened response of eosinophilic asthmatic patients to the CRTH2 antagonist OC000459. Allergy 69, 1223–1232 (2014).

26. Gonem, S. et al. Fevipiprant, a prostaglandin D2 receptor 2 antagonist, in patients with persistent eosinophilic asthma: a single-centre, randomised, double-blind, parallel-group, placebo-controlled trial. Lancet Respir Med. 4, 699–707 (2016).

27. Xue, L. et al. Prostaglandin D2 and leukotriene E4 synergize to stimulate diverse TH2 functions and TH2 cell/neutrophil crosstalk. J Allergy Clin Immunol. 135, 1358–1366 e1351–1311 (2015).

28. Salimi, M. et al. Cysteinyl leukotriene E4 activates human group 2 innate lymphoid cells and enhances the effect of prostaglandin D2 and epithelial cytokines. J Allergy Clin Immunol. 140, 1090–1100 (2017).

29. Hinks, T. S. et al. Innate and adaptive T cells in asthmatic patients: Relationship to severity and disease mechanisms. J Allergy Clin Immunol. 136, 323–333 (2015).

30. Xue, L. et al. Leukotriene E4 activates human Th2 cells for exaggerated proinflammatory cytokine production in response to prostaglandin D2. J Immunol. 188, 694–702 (2012).

31. Lamkhioued, B. et al. Increased expression of eotaxin in bronchoalveolar lavage and airways of asthmatics contributes to the chemotaxis of eosinophils to the site of inflammation. J Immunol. 159, 4593–4601 (1997).

32. Lilly, C. M. et al. Expression of eotaxin by human lung epithelial cells: induction by cytokines and inhibition by glucocorticoids. J Clin Invest. 99, 1767–1773 (1997).

33. Palikhe, N. S. et al. Elevated levels of circulating CD4(+) CRTh2(+) T cells characterize severe asthma. Clin Exp Allergy 46, 825–836 (2016).

34. Stoeckle, C..& Simon, H. U. CD8(+) T cells producing IL-3 and IL-5 in non-IgE-mediated eosinophilic diseases. Allergy 68, 1622–1625 (2013).

35. Tsitsiou, E. et al. Transcriptome analysis shows activation of circulating CD8+ T cells in patients with severe asthma. J Allergy Clin Immunol. 129, 95–103 (2012).

36. Kuo, C. S. et al. A Transcriptome-driven Analysis of Epithelial Brushings and Bronchial Biopsies to Define Asthma Phenotypes in U-BIOPRED. Am J Respir Crit Care Med. 195, 443–455 (2017).

37. Koya, T. et al. CD8+ T cell-mediated airway hyperresponsiveness and inflammation is dependent on CD4+IL-4+ T cells. J Immunol. 179, 2787–96 (2007).

38. Miyahara, N. et al. Effector CD8+ T cells mediate inflammation and airway hyper-responsiveness. Nat Med. 10, 865–869 (2004).

39. Schedel, M. et al. 1,25D3 prevents CD8(+)Tc2 skewing and asthma development through VDR binding changes to the Cyp11a1 promoter. Nat Commun. 7, 10213 (2016).

40. Pavord, I. D. et al. Mepolizumab for severe eosinophilic asthma (DREAM): a multicentre, double-blind, placebo-controlled trial. Lancet 380, 651–659 (2012).

41. Kay, A. B., Phipps, S. & Robinson, D. S. A role for eosinophils in airway remodelling in asthma. Trends Immunol. 25, 477–482 (2004).

42. Frigas, E., Motojima, S. & Gleich, G. J. The eosinophilic injury to the mucosa of the airways in the pathogenesis of bronchial asthma. Eur Respir J Suppl. 13, 123s–135s (1991).

43. Green, R. H. et al. Asthma exacerbations and sputum eosinophil counts: a randomised controlled trial. Lancet 360, 1715–1721 (2002).

44. Dent, G. et al. Contribution of eotaxin-1 to eosinophil chemotactic activity of moderate and severe asthmatic sputum. Am J Respir Crit Care Med. 169, 1110–1117 (2004).

45. Schwarze, J. et al. Transfer of the enhancing effect of respiratory syncytial virus infection on subsequent allergic airway sensitization by T lymphocytes. J Immunol. 163, 5729–5734 (1999).

46. Sampson, S. E., Sampson, A.P. & Costello, J. F. Effect of inhaled prostaglandin D2 in normal and atopic subjects, and of pretreatment with leukotriene D4. Thorax 52, 513–518 (1997).

47. Mastalerz, L. et al. Induced sputum supernatant bioactive lipid mediators can identify subtypes of asthma. Clin Exp Allergy 45, 1779–1789 (2015).

48. Takamoto, M..& Sugane, K. Synergism of IL-3, IL-5, and GM-CSF on eosinophil differentiation and its application for an assay of murine IL-5 as an eosinophil differentiation factor. Immunol Lett. 45, 43–46 (1995).

49. Choi, J., Callaway, Z., Kim, H. B., Fujisawa, T. & Kim, C. K. The role of TNF-alpha in eosinophilic inflammation associated with RSV bronchiolitis. Pediatr Allergy Immunol. 21, 474–479 (2010).

50. Kim, M. R. et al. Transgenic overexpression of human IL-17E results in eosinophilia, B-lymphocyte hyperplasia, and altered antibody production. Blood 100, 2330–2340 (2002).

51. Oliveira, S. H. et al. Increased responsiveness of murine eosinophils to MIP-1beta (CCL4) and TCA-3 (CCL1) is mediated by their specific receptors, CCR5 and CCR8. J Leukoc Biol. 71, 1019–1025 (2002).

52. Hanania, N. A. et al. Efficacy and safety of lebrikizumab in patients with uncontrolled asthma (LAVOLTA I and LAVOLTA II): replicate, phase 3, randomised, double-blind, placebo-controlled trials. Lancet Respir Med. 4, 781–796 (2016).

53. Ritchie, M. E. et al. Limma powers differential expression analyses for RNA-sequencing and microarray studies. Nucleic Acids Research 43, e47 (2015).

